# Targeting Scavenger Receptor Type B-1 (SR-B1) and Cholesterol Inhibits Entry of SARS-CoV-2 Pseudovirus in Cell Culture

**DOI:** 10.1101/2020.12.14.420133

**Authors:** Stephen E. Henrich, Kaylin M. McMahon, Nicole Palacio, Pankaj Bhalla, Pablo Penaloza-MacMaster, C. Shad Thaxton

**Affiliations:** Department of Urology, Feinberg School of Medicine, Northwestern University, 303 E. Chicago Ave., Chicago, Illinois, USA; Department of Microbiology-Immunology, Feinberg School of Medicine, Northwestern University, 303 E. Chicago Ave., Chicago, Illinois, USA; Department of Dermatology, Feinberg School of Medicine, Northwestern University, 303 E. Chicago Ave., Chicago, Illinois, USA; Simpson Querrey Institute for BioNanotechnology, Northwestern University, 303 E. Chicago Ave., Chicago, Illinois, USA; Robert H. Lurie Comprehensive Cancer Center, Northwestern University, 303 E. Chicago Ave., Chicago, Illinois, USA

## Abstract

The novel human coronavirus, severe acute respiratory syndrome coronavirus 2 (SARS-CoV-2), emerged in Wuhan, China in late 2019 and has now caused a global pandemic. The disease caused by SARS-CoV-2 is known as COVID-19. To date, few treatments for COVID-19 have proven effective, and the current standard of care is primarily supportive. As a result, novel therapeutic strategies are in high demand. Viral entry into target cells is frequently sensitive to cell membrane lipid composition and membrane organization. Evidence suggests that cell entry of SARS-CoV-2 is most efficient when the target cell plasma membrane is replete with cholesterol; and recent data implicate cholesterol flux through the high-affinity receptor for cholesterol-rich high-density lipoprotein (HDL), called scavenger receptor type B-1 (SR-B1), as critical for SARS-CoV-2 entry. Here, we demonstrate that a cholesterol-poor synthetic biologic high-density lipoprotein (HDL NP) targets SR-B1 and inhibits cell entry of a SARS-CoV-2 spike protein pseudovirus. Human cells expressing SR-B1 are susceptible to SARS-CoV-2 infection, and viral entry can be inhibited by 50-80% using HDL NPs in an SR-B1-dependent manner. These results indicate that HDL NP targeting of SR-B1 is a powerful potential therapy to combat COVID-19 and other viral diseases.

## Introduction

COVID-19 disparately impacts the aging population, with the median age of COVID-19-related death being 78 years.^1^ Individuals with pre-existing conditions such as coronary artery disease,^2-3^ hypertension,^4^ and diabetes^5^ are also disproportionately susceptible to serious sequalae from severe acute respiratory syndrome coronavirus 2 (SARS-CoV-2) infection. However, the precise reasons why advanced age and pre-existing conditions lead to these disparities remain unclear. Prior evidence indicates that SARS-CoV-1 infectivity is highly sensitive to cholesterol levels in the host cell.^6^ Indeed, several recent studies demonstrate that this is also the case for SARS-CoV-2.^7-9^ Specifically, cholesterol-rich membrane domains constitute viral entry points for SARS-CoV-2.^7^ Moreover, cholesterol helps traffic angiotensin converting enzyme 2 (ACE2) to these domains and increases the binding affinity for SARS-CoV-2 to the cell surface.^7^ Conversely, reducing cellular cholesterol reduces viral infectivity. These findings provide valuable mechanistic insight into the existing disparities in COVID-19 outcomes for the elderly and those with pre-existing cardiovascular conditions and suggest that targeted reduction of cellular cholesterol may be a viable therapeutic strategy. Here, we sought to determine whether a targeted, cholesterol-reducing nanoparticle agent could inhibit SARS-CoV-2 entry.

Coronaviruses are enveloped viruses with positive stranded RNA genomes. Enveloped viruses typically infect their host cells in a two-step process whereby viral surface proteins first bind to receptors on the cell surface to initiate viral attachment, followed by a fusion event which leads to internalization of the virion by the host cell. Each virus in the coronavirus family expresses three structural proteins that are incorporated into the viral capsid, the membrane (M), envelope (E), and spike (S) proteins. The S protein is most important for viral attachment to host cells. For SARS-CoV-1 and 2, viral entry is facilitated by binding of the S protein to its receptor on the surface of certain host cells, ACE2. Furthermore, the high-affinity receptor for native cholesterol-rich high-density lipoproteins (HDL), called scavenger receptor type B-1 (SR-B1), has recently been implicated as a co-receptor to facilitate the entry of SARS-CoV-2, potentially in lock-step with HDL, into cells of the host airway.^10^ SR-B1 is expressed by hepatocytes,^11-12^ immune cells^12-14^ and type II pneumocytes,^15^ the latter representing a major cell type targeted by SARS-CoV-2. SR-B1 has previously been appreciated as a co-receptor for other viruses and pathogens, including hepatitis C virus^16^ and Plasmodium species.^17^

Despite this basic knowledge regarding the mechanism by which SARS-CoV-2 infects host cells, knowledge regarding therapeutic targets that could be exploited to inhibit SARS-CoV-2 infectivity remains limited. Most work has been focused on antibodies or ACE2 receptor decoys that target viral antigens, like the S protein, to reduce the productive interactions between the virus and host cells.^18-21^ However, strategies that target host cells to prevent SARS-CoV-2 entry are less well studied. It is known that SARS-CoV-1 and SARS-CoV-2 utilize cholesterol and monosialotetrahexosylganglioside 1 (GM1)-rich lipid microdomains, or lipid rafts, for viral entry in cultured mammalian cells. Specifically, treating cells with the non-specific cholesterol sequestrant, methyl-β-cyclodextrin (MβCD), prior to introducing SARS-CoV-1 or SARS-CoV-2, leads to significantly reduced infection. Furthermore, recent data demonstrate that modulating cell cholesterol biosynthesis may be a critical pathway that controls SARS-CoV-2 entry into host cells.^22^ Taken together, a significant amount of data suggest that the entry of SARS-CoV-2 into host cells is cholesterol-dependent and that targeting cell entry with an agent that modulates cell membrane and cellular cholesterol metabolism may provide a unique therapy to prevent SARS-CoV-2 entry into host cells.

HDLs are multi-functional particles that greatly impact inflammation and are taking on increasing importance as guardians of the integrity of endothelial and epithelial barrier function throughout the body. At the most basic level, HDLs are dynamic nanoscale particles (7-13 nm in diameter) that circulate in the bloodstream of mammals and transport cholesterol. HDLs are specifically notable for their functions in so-called reverse cholesterol transport, whereby they remove cholesterol from peripheral cells such as macrophages and deliver cholesterol back to hepatocytes for excretion in the bile. As a result, HDLs can potently modulate cell cholesterol metabolism. Our group and others have developed and tested synthetic HDLs for a variety of therapeutic^23-27^ and imaging^28-29^ purposes. In one iteration, an inorganic core nanoparticle is used as a template to assemble the protein (apolipoprotein A-I) and lipids naturally associated with the surface of native cholesterol-rich HDLs. By virtue of their surface mimicry, HDL NPs tightly bind SR-B1. However, because synthetic HDLs lack a core of esterified cholesterol, and other lipids, they have been shown to differentially modulate cell membrane cholesterol and, by way of reducing cell cholesterol, stimulate a compensatory increase in cellular cholesterol biosynthesis. From a therapeutic perspective, HDL NPs modulate cell membrane cholesterol and lipid rafts, which has been shown to potently inhibit the uptake of extracellular lipid vesicles, often referred to as exosomes, which share many properties with viruses.^30-31^ In addition, HDL NP modulation of cell cholesterol has been leveraged to develop potent anticancer therapies for tumor types that heavily depend upon cholesterol uptake through SR-B1.^26^ And, finally, tuning the lipids bound to the outer surface of HDL NP enables profound targeting of Gram-negative bacterial lipopolysaccharide (LPS) to attenuate the activation of NF-κB and attendant cytokine expression.^32^ Because of these properties, and a growing body of evidence demonstrating the dependence of SARS-CoV-2 infection upon cell cholesterol, we employed synthetic HDLs to determine whether targeted reduction of cellular cholesterol would inhibit viral entry of a SARS-CoV-2 spike protein pseudovirus.

## Materials and Methods

### Cell culture

HEK293 cells expressing the human ACE2 receptor and HepG2 cells were cultured in DMEM containing 10% fetal bovine serum (FBS) and 1% penicillin/streptomycin (PenStrep). All cell cultures were maintained at 37°C and 5% CO_2_. Frozen aliquots of 10^6^ cells were used and cells were passaged fewer than 10 times for all experiments.

### High-density lipoprotein nanoparticle (HDL NP) synthesis

HDL NPs were synthesized according to previously published protocols. Briefly, particle synthesis was initiated by adding apoA-1 (MyBioSource) at a 5:1 mole ratio to a colloidal suspension of 5 nm diameter citrate-stabilized gold nanoparticles (AuNP) (Ted Pella). The suspension was vortexed briefly and then incubated at RT for 1 h on a flat-bottom shaker. Next, a phospholipid--1,2-dipalmitoyl-*sn*-glycero-3-phosphoethanolamine-*N*-[3-(2-pyridyldithio)propionate] (PDP PE) (Avanti Polar Lipids)—was added to the suspension at a 250-fold molar excess to [AuNP], followed by two additional phospholipids, 1,2-dilinoleoyl-*sn*-glycero-3-phospho-(1’-rac-glycerol) (18:2 PG) and cardiolipin (Avanti Polar Lipids), each at 125-fold molar excess to the AuNP. The lipids were incubated with the suspension in a mixture of ethanol and water (1:4) for 4 h at RT with gentle mixing on a flat-bottom shaker. The HDL NPs were then purified by tangential flow filtration. The HDL NP concentration was determined by UV-Vis spectroscopy and Beer’s law (ε_AuNP_ = 9.696 × 10^6^ M^-1^cm^-1^, λ_max_ = 520 nm). Particle hydrodynamic diameter was determined by dynamic light scattering.

### SARS-CoV-2 pseudovirus production

First, 293T cells grown to ∼70-80% confluency. Before transfection, the culture media was aspirated and changed to serum-free DMEM. Cells were transfected with a SARS-CoV-2 spike expression plasmid (NR-52310 from BEI Resources) using PEI. After 24 hours, VSV-dG*G-GFP virus was added at an MOI of 3, and incubated for 2 hours, rocking gently every 15 min. Two hours after infection, the media was replaced with DMEM with 2% FBS and then cultured for another 48 hours. Finally, the cell supernatant was removed and filtered through a 0.45 µM filter and then concentrated using an Amicon concentrator (UFC910024 from Sigma Millipore).

### Viral entry inhibition experiments

HepG2 and HEK293 (ACE2) cells were seeded in 24-well or 96-well tissue culture plates at 50,000 or 10,000 cells/ well, respectively. After 24 h, cells were washed once in PBS and SARS-CoV-2 pseudovirus was added to the cells in culture medium containing 1% FBS at a concentration of 350 fluorescence-forming units (FFU)/mL. For indicated groups, HDL NPs were then added to the wells at a final concentration of 50 nM. Plates were then placed in an IncuCyte S3 to enable live cell imaging. Phase and green fluorescence images were acquired every hour for 48 h. IncuCyte S3 analysis software was used to process the data. For experiments using SR-B1 siRNA or scramble control, the same protocol was used as above with the exception that 24 h after cell seeding, cells were treated with SR-B1 siRNA or scramble RNA using Lipofectamine RNAiMAX according to the manufacturer’s instructions for 48 h prior to addition of virus with or without HDL NP.

### SR-B1 knockdown experiments

HEK293 (ACE2) cells were plated at 400,000 cells per well in 6-well tissue culture plates. When cells had reached approximately 70% confluence, siRNA targeting SR-B1 (Ambion) and negative control scramble RNA (Ambion) were prepared using Lipofectamine RNAiMAX in Opti-MEM according to the manufacturer’s instructions. Pre-prepared RNA was added to the cells at 30 nM and SR-B1 knockdown proceeded for 48 h. Cell lysates were harvested using M-PER lysis buffer, samples were centrifuged for 10 min at 14,000 x *g* to pellet cellular debris. The supernatant was then transferred to a new tube and protease and phosphatase inhibitors were added prior to processing for western blot.

### Western blot

Sample protein concentration was determined using bicinchoninic acid (BCA) assay. Samples were then normalized to total protein, mixed with 4X Laemmli loading buffer containing β-mercaptoethanol, and boiled for 10 minutes at 100 °C. Proteins were resolved using a 4%-20% polyacrylamide gel (120 V, 1 h) and transferred to a 0.45 μm PVDF membrane (60 V, 1 h). The membrane was blocked using 5% milk in Tris buffered saline (TBS) and Tween-20 (0.1%) for 1 hour. SR-B1 antibody was applied (1:2000) (Abcam, ab52629) and incubated overnight at 4 °C. Blot was washed 10 minutes (3X) in TBST (0.1% Tween-20) secondary goat anti-rabbit antibody (BioRad, 1721019) was applied (1:1000) for 1 hour at R.T. and blot was washed (3X) same as described above. Protein was detected using enhanced chemiluminescence (ECL) detection (Bio-Rad, 1705060) and an Azure 300 (Azure Biosystems) gel imaging system.

## Results and Discussion

Synthetic HDL NPs target the high-affinity receptor for native cholesterol-rich HDLs, SR-B1. As such, we assayed for the presence of SR-B1 in model cell lines of SARS-CoV-2 infection. The HepG2 and HEK293 (ACE2 over-expressing) cell lines both expressed SR-B1 as determined by Western blot (Figure 1 a,b). Moreover, treating HEK293 (ACE2) and HepG2 cells with siRNA against SR-B1 was shown to knock down SR-B1 protein expression (Figure 1 a,b) in each of these cell lines. The latter result is important for downstream SR-B1-dependence studies.

**Figure 1.**
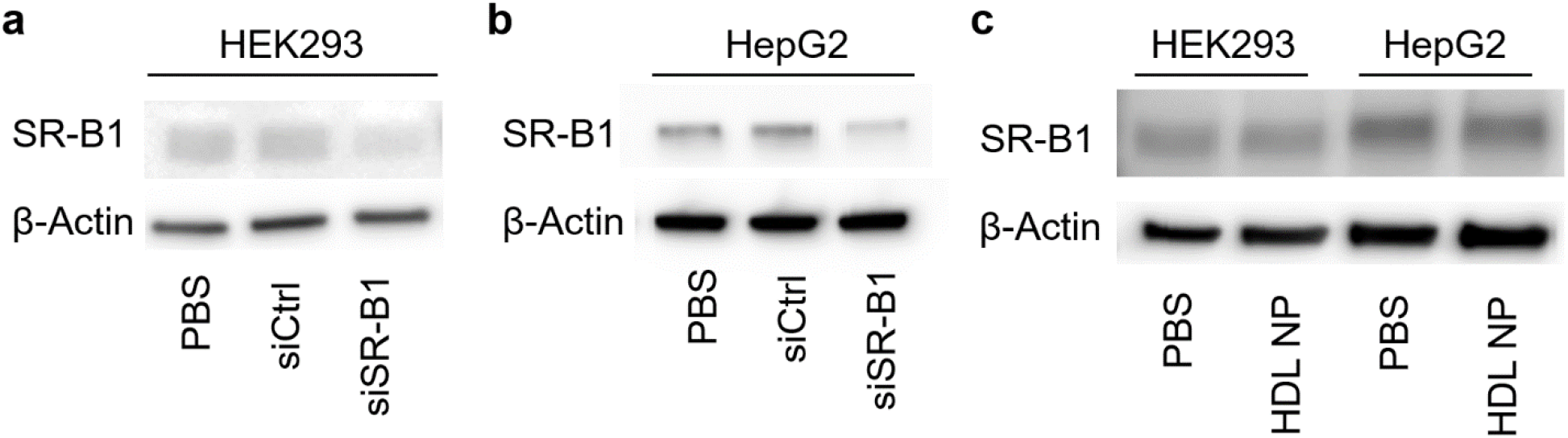
a-b) HEK293 (ACE2 over-expressing) and HepG2 cells express the native receptor for mature spherical HDL, SR-B1. c) HDL NP treatment does not alter SR-B1 expression.

Next, we treated HEK293 (ACE2) and HepG2 cells with PBS control or HDL NPs to determine any alterations in SR-B1 expression. We found that HDL NPs did not alter SR-B1 expression in these cell lines (Figure 1 c) at the concentration and treatment duration used for viral entry experiments later in the study (50 nM HDL NP for 48 h).

Having established that HEK293 (ACE2) and HepG2 cells express SR-B1 and would therefore potentially be susceptible to HDL NP targeting, we next performed experiments to determine whether HDL NP treatment had any impact on the infectivity of a GFP-expressing SARS-CoV-2 spike protein pseudovirus (SARS-CoV-2 pseudovirus). Cells were treated with HDL NPs (50 nM) at the same time as SARS-CoV-2 pseudovirus was introduced, and cells were monitored with live cell fluorescence imaging for 48 h for GFP expression. Live cell imaging snapshots (Figure 2a, 3a) and quantifications (Figure 2b, 3b) showed that HDL NP treatment significantly inhibited SARS-CoV-2 pseudovirus infectivity by approximately 55% in HEK293 (ACE2) cells and 80% in HepG2 cells after 48 h. These results are consistent with the relative SR-B1 expression of the two cell lines, with HepG2 cells expressing more SR-B1 than HEK293 (ACE2) cells (Figure 1).

**Figure 2.**
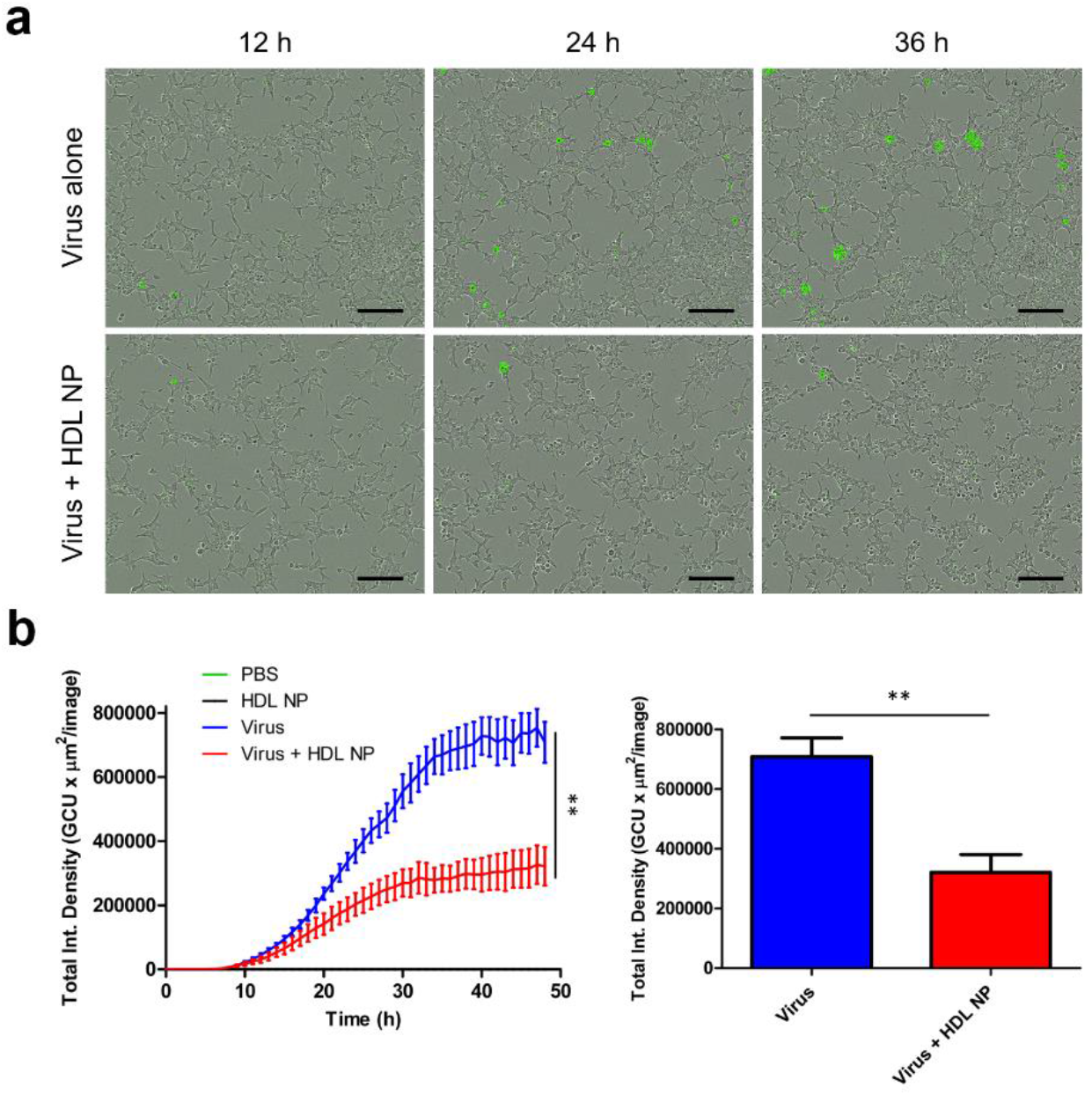
HDL NP treatment inhibits infection of SARS-CoV-2 pseudovirus in HEK293 (ACE2) cells. a) Live cell imaging snapshots of HEK293 (ACE2) cells treated with GFP-expressing SARS-CoV-2 pseudovirus with or without HDL NP (50 nM) co-treatment. b) Quantification of GFP^+^ HEK293 (ACE2) cells (total integrated green fluorescence density) after 48 h of infection with or without HDL NP co-treatment. Scale bars = 200 µm.

**Figure 3.**
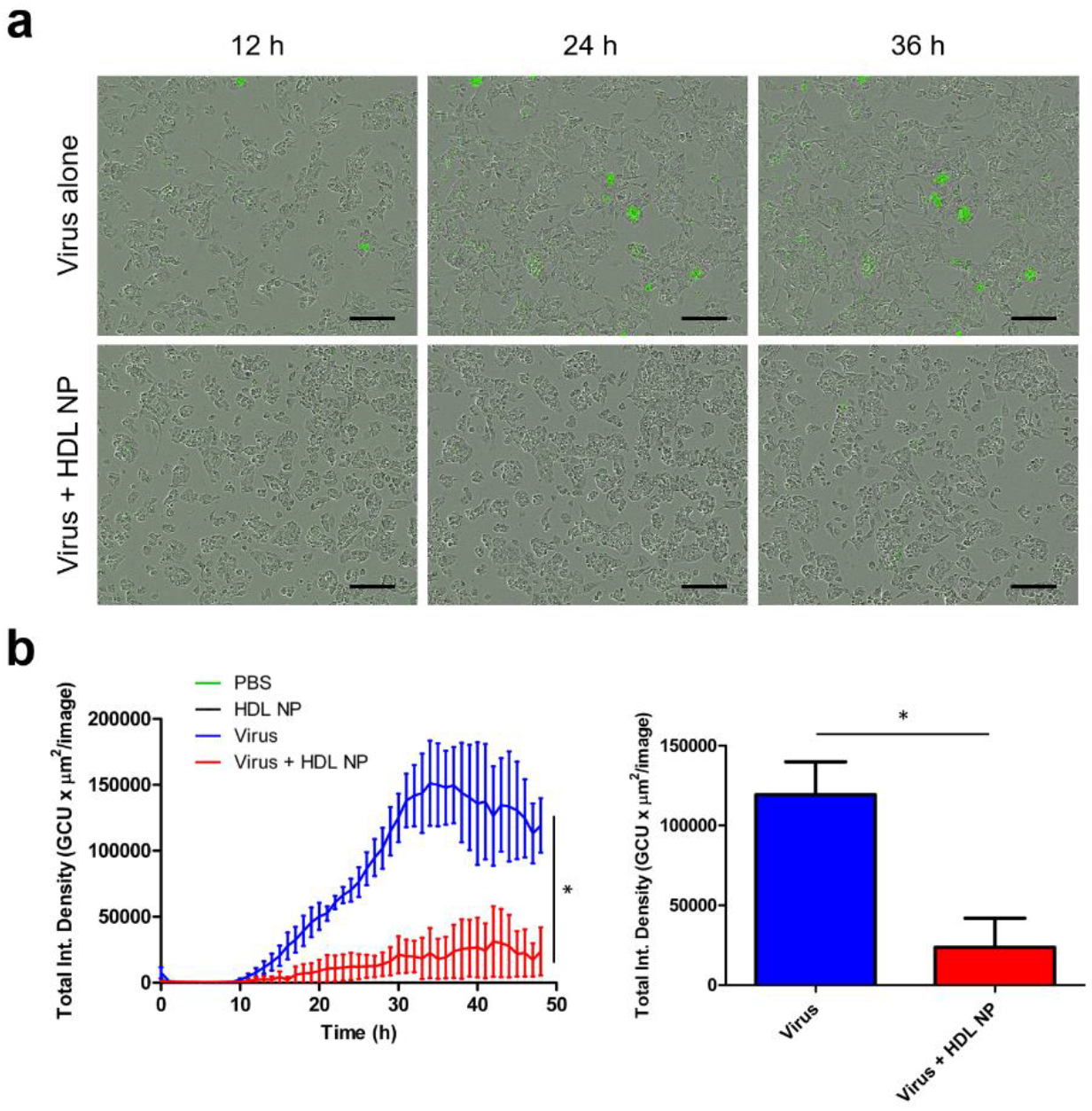
HDL NPs inhibit infection of SARS-CoV-2 pseudovirus in HepG2 cells. a) Live cell imaging snapshots of HepG2 cells treated with GFP-expressing SARS-CoV-2 pseudovirus with or without HDL NP (50 nM) co-treatment. b) Quantification of GFP^+^ HepG2 cells (total integrated green fluorescence density) after 48 h of infection with or without HDL NP co-treatment. Scale bars = 200 µm.

With the knowledge the SR-B1 has been identified as a possible co-receptor for SARS-CoV-2, we then investigated whether knocking down SR-B1 had any effect on SARS-CoV-2 infectivity. To do this, we transfected HEK293 (ACE2) cells for 48 h with anti-SR-B1 siRNA or a scrambled RNA control using Lipofectamine RNAiMAX, and then introduced the SARS-CoV-2 pseudovirus and allowed infection to proceed for 48 h. We found that knocking down SR-B1 led to reduced viral entry (Figure 4a, top two rows; Figure 4b), consistent with the hypothesis that SR-B1 is a co-receptor for SARS-CoV-2.

**Figure 4.**
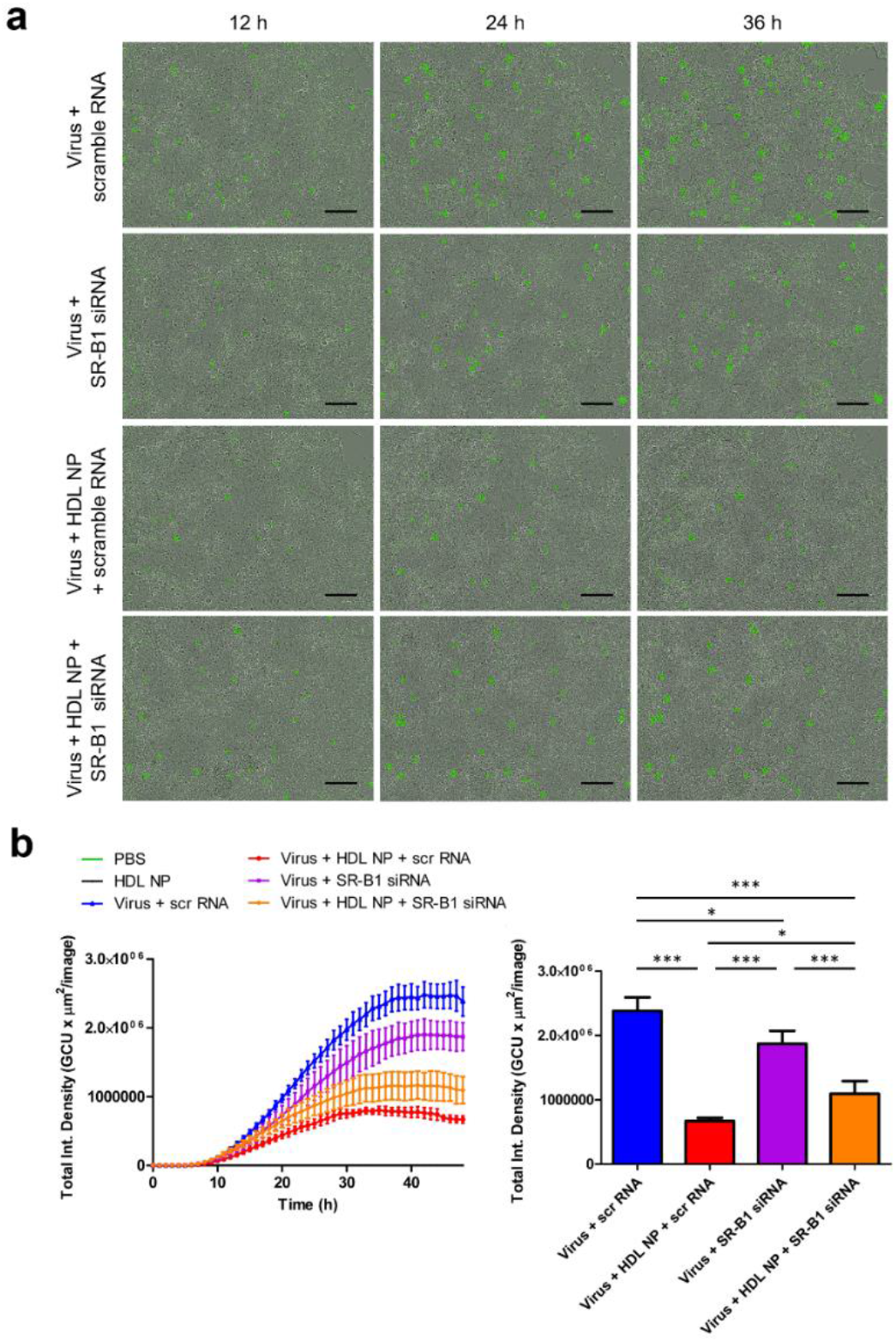
a) Live cell imaging of HEK293 (ACE2) cells infected with SARS-CoV-2 pseudovirus subjected to treatment regimens of HDL NPs, anti-SR-B1 siRNA, and/or scramble RNA. b) Quantification of GFP^+^ cells (total integrated green fluorescence density) in groups infected with SARS-CoV-2 pseudovirus with or without treatment with HDL NPs, SR-B1 siRNA, or scramble RNA control. Scale bars = 200 µm.

Finally, we sought to interrogate the complex relationship between viral entry, HDL NP inhibition, and SR-B1 by knocking down SR-B1 prior to infection, in conjunction with HDL NP treatment. Viral entry experiments were performed using SARS-CoV-2 pseudovirus in HEK293 (ACE2) cells and HepG2 cells treated with anti-SR-B1 siRNA for 48 h to reduce the expression of SR-B1. Subsequently, cells were treated with SARS-CoV-2 pseudovirus with or without HDL NP (50 nM) for an additional 48 h. Live cell imaging revealed that cells treated with HDL NPs in addition to SR-B1 knockdown exhibited lower rates of viral entry than merely reducing SR-B1 alone (Figure 4); however, the rate of viral entry was not as low as virus + HDL NP without siRNA (Figure 4).

Taken together, these results demonstrate that SARS-CoV-2 viral entry is reduced upon SR-B1 knockdown, suggesting that SR-B1 is a co-receptor for SARS-CoV-2; moreover, data support that HDL NPs target SR-B1 to inhibit SARS-CoV-2 entry.

## Conclusion

The novel SARS-CoV-2 virus has caused a global pandemic. Recent data clearly demonstrate that cellular infection by SARS-CoV-2 is impacted by cell membrane and cellular cholesterol levels, as well as requiring the presence of the putative co-receptor SR-B1. As such, we employed our synthetic HDL NP to target SR-B1 on the surface of cultured cells known to be targets of SARS-CoV-2. Our data clearly demonstrate potent reduction of viral entry into target cells at low nM therapeutic doses of the HDL NP. Furthermore, inhibition of SARS-CoV-2 is long lasting, up to 48 hours after the initiation of culture. No untoward side effects of HDL NP therapy were observed. Furthermore, using siRNA to reduce the expression of SR-B1 in the host cell types, data show that the HDL NPs are actively targeting this receptor; and data support that SR-B1 is a co-receptor for SARS-CoV-2 as reduced viral entry is observed after SR-B1 knockdown in the absence of HDL NP treatment. Overall, HDL NPs are a promising therapeutic candidate to inhibit SARS-CoV-2 entry and are also a candidate therapy for the general reduction of viral infection of host cells that express SR-B1 and depend upon cell membrane cholesterol and lipid raft integrity for cell entry.

## Acknowledgements

S.E.H. gratefully acknowledges support from an NRSA F30 fellowship from the NIH (CA225133-02). The authors thank the Analytical Bionanotechnology Equipment Core (ANTEC) and Skin Biology & Disease Resource-based Center (SBDRC) facilities at Northwestern University.

